# Generalized Multi-SNP Mediation Intersection-Union Test

**DOI:** 10.1101/780767

**Authors:** Wujuan Zhong, Toni Darville, Xiaojing Zheng, Jason Fine, Yun Li

## Abstract

To elucidate the molecular mechanisms underlying genetic variants identified from genome-wide association studies (GWAS) for a variety of phenotypic traits encompassing binary, continuous, count, and survival outcomes, we propose a novel and flexible method to test for mediation that can simultaneously accommodate multiple genetic variants and different types of outcome variables. Specifically, we employ the intersection-union test approach combined with likelihood ratio test to detect mediation effect of multiple genetic variants via some mediator (for example, the expression of a neighboring gene) on outcome. We fit high-dimensional generalized linear mixed models under the mediation framework, separately under the null and alternative hypothesis. We leverage Laplace approximation to compute the marginal likelihood of outcome and use coordinate descent algorithm to estimate corresponding parameters. Our extensive simulations demonstrate the validity of our proposed method and substantial, up to 97%, power gains over alternative methods. Applications to real data for the study of *Chlamydia trachomatis* infection further showcase advantages of our method. We believe our proposed method will be of value and general interest in this post-GWAS era to disentangle the potential causal mechanism from DNA to phenotype for new drug discovery and personalized medicine.

## 1. Introduction

Mediation analysis studies how the mediator variable transmits the independent variable’s effect on the outcome (MacKinnon et al., 2007). Most mediation studies focus on outcomes following Gaussian distribution. Non-Gaussian outcomes, such as binary, count and time-to-event responses (e.g. disease status, time until death), are commonly present in research but have been under-studied. In mediation analysis, non-Gaussian outcomes from the exponential family distribution can be properly handled by generalized linear models (GLM) and time-to-event outcomes can be accommodated using a proportional hazards Cox model (Preacher, 2015). For example, (O’Rourke and Vazquez, 2019) discusses challenges in mediation analysis of zero-inflated count outcomes and describes how to fit Poisson or negative binomial models and (Cheng et al., 2018) attempts to decompose the direct, mediation and total effects for zero-inflated count outcomes from a causal inference perspective.

Generalized linear mixed models (GLMM) (McCullagh and Nelder, 1989; McCulloch and Searle, 2001; McCulloch et al., 2008) are an extension of GLM where random effects are accommodated among the predictors. GLMM are commonly be applied to data where observations are not independent, for instance in studies with repeated measures. In genetics and genomics studies, GLMM is widely used to test associations between non-Gaussian traits and a set of genetic variants (Yan et al., 2015; Chen et al., 2016, 2019; Park et al., 2018) when genetic relationship among study subjects needs to be taken into account. Similarly for survival outcome, mixed effects Cox models (Vaida and Xu, 2000; Pankratz et al., 2005) have been developed as an extension of proportional hazards Cox model to allow explicitly modeling of random effects.

Likelihood-based inference for GLMM can be difficult, because it usually involves high-dimensional integrals (McCulloch et al., 2008). For this reason, various strategies have been proposed to approximate the likelihood function for GLMM, including Laplace approximation (Raudenbush et al., 2000), penalized quasi-likelihood (PQL) (Breslow and Clayton, 1993), and Markov chain Monte Carlo (MCMC) algorithms (Gilks, 1996). An excellent review paper about GLMM in practice exists (Bolker et al., 2009). For time-to-event outcome, Laplace approximation has been applied to approximate likelihood function for mixed effects Cox models (Pankratz et al., 2005). To maximize the approximated likelihood function, coordinate descent (Fu, 1998; Daubechies et al., 2004) is broadly used, such as for GLM with elastic net (Friedman et al., 2010), graphical Lasso (Friedman et al., 2008) and GLMM with Lasso (Schelldorfer et al., 2014). Coordinate descent is simple and convenient to employ and can achieve satisfactory performance when carefully implemented.

Mediation analysis was firstly proposed by Baron and Kenny to study the association between an independent variable and an outcome by adding an intermediate variable, which is called the mediator (Baron and Kenny, 1986). In genetics and genomics studies, researchers are interested in testing mediation effects of the genetic variant(s), mostly single nucleotide polymorphisms (SNPs) on the outcome through certain mediator (e.g., the expression level of a neighboring gene). Baron and Kenny’s classic mediation approach has been extended to accommodate high-dimensional mediators (Huang and Pan, 2016; Zhang et al., 2016). Huang et al.’s methods are kernel-based regression methods and use variance component score statistic to test for mediation but these methods assume a priori known expression quantitative trait loci (eQTLs) (Huang et al., 2015, 2016). To address lack of knowledge regarding eQTLs, we have extended Baron and Kenny’s framework to handle mediation effect of high-dimensional genetic variants on a continuous outcome (Zhong et al., 2019). To the best of our knowledge, none of the existing methods can jointly test mediation effects of multiple correlated SNPs on a non-Gaussian outcome. We propose a generalized multi-SNP mediation intersection-union test to accommodate both mediation and direct effects of multiple correlated SNPs on non-Gaussian outcomes without a prior knowledge of eQTLs. Similar to our previously developed SMUT method (Zhong et al., 2019), the method proposed in this work is an extension of Baron and Kenny’s framework and leverages intersection-union test (IUT) to decompose mediation into two separate regression models. Our proposed method SMUT_GLM and SMUT_PH deals with two categories of non-Gaussian outcomes. SMUT_GLM handles an outcome from an exponential family distribution by fitting a generalized linear mixed model and SMUT_PH accommodates a survival by fitting a mixed effects Cox proportional hazards model.

The rest of this article is organized as follows. In Section 2, we present details of our proposed SMUT_GLM and SMUT_PH methods, followed by simulation studies and real data application in Section 3 and Section 4, respectively. Finally, Section 5 concludes the article with some discussions.

## 2. Methods

### 2.1 Notation

Without loss of generality, we assume that we have four types of data, namely, genotypes (as the potential causal variables), gene expression measurements (as the mediator, which can be other types of molecular measures such as metabolite levels or protein abundances), phenotypic trait (as the final outcome) and other covariates (e.g. age, gender). Let *G* = (*G*_1_, *G*_2_, …, *G*_*q*_) be the *n* by *q* genotype matrix, where *n* is sample size, *q* is the total number of genetic markers, and *G*_*j*_ = (*G*_1*j*_, *G*_2*j*_, …, *G*_*nj*_)^*T*^ is the vector of genotypes for the samples at marker *j*, *j* = 1, 2, …, *q*. We consider an additive model with *G*_*ij*_ taking values 0,1,2, measuring the number of copies of the minor allele. Let *X*_*ij*_ denote the *j*th covariate variable (e.g. age, gender) for the *i*th individual, *i* = 1, 2, …, *n*; *j* = 1, 2, …, *p*.

### 2.2 SMUT_GLM and SMUT_PH model

SMUT_GLM and SMUT_PH model the effects of SNPs on the outcome mediated by the expression level of a single gene via two models, namely a mediator model and an outcome model. We assume the expression level is continuous and consider a linear model for the mediator model (equation 1). As for the outcome model, we fit GLMM if the outcome random variable follows an exponential family distribution (equation 2); we fit mixed effects proportional hazards Cox model if the outcome is time-to-event (equation 3). Let *Y*_*i*_ denote the outcome for the *i*th individual. For survival outcome, *Y*_*i*_ = (*T*_*i*_, *δ*_*i*_) includes the time *T*_*i*_ = *min*(*Z*_*i*_, *C*_*i*_), where *Z*_*i*_ is the time to the event of interest and *C*_*i*_ is the censoring time, and the censoring status *δ*_*i*_; *δ*_*i*_ = 1 indicates the occurrence of the given event and *T*_*i*_ is the survival time; *δ*_*i*_ = 0 indicates a censored sample.

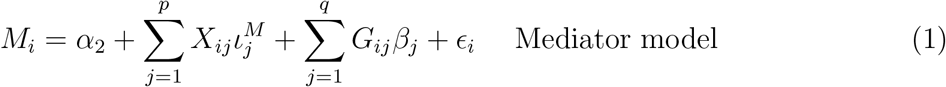

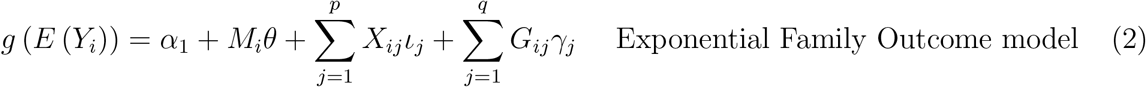

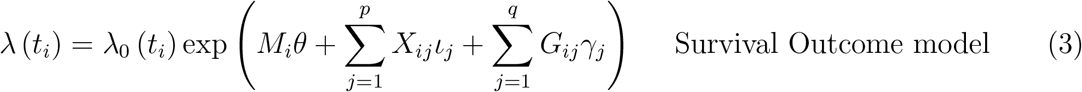

where *i* = 1, 2, …, *n* indexes the *n* individuals; *q* is the number of SNPs; *ϵ*_*i*_ ∼_*i.i.d.*_ *N*(0, 1), *i* = 1, 2, …, *n*; *g* is the link function in GLM. Here 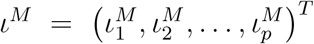 and 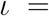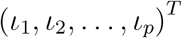 are coefficient vectors for the *p* covariates in the mediator and outcome model, respectively; *β* = (*β*_1_, *β*_2_, …, *β*_*q*_)^*T*^ is the SNP effect on the mediator *M*; *θ* is the mediator effect on the outcome; *βθ* is the mediation effect of the SNPs via mediator *M*; *γ* = (*γ*_1_, *γ*_2_, …, *γ*_*q*_)^*T*^ includes the direct effects of the *q* SNPs and mediation effects via mediators other than *M*. For presentation brevity, we will use direct effects to refer to the aggregated effects including SNPs’ direct effects and mediation effects via other mediators.

Following our previously developed SMUT method (Zhong et al., 2019), we employ intersection-union test (IUT) (Berger and Hsu, 1996) to decompose the hypothesis testing of the mediation effect *βθ* into two sub-hypotheses: 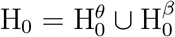 and 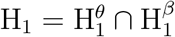, where 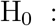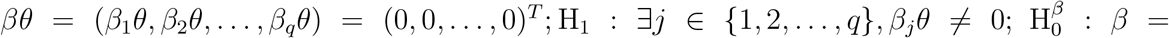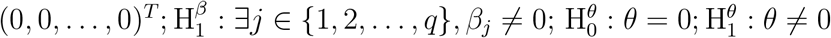.

Suppose the *p* value for testing *β* being zero is *p*_1_; and the *p* value for testing *θ* being zero is *p*_2_. Then the *p* value for testing *βθ* being zero, using IUT, is the maximum of *p*_1_ and *p*_2_. In the following sections, we provide details regarding how to separately test *β* and *θ* to obtain *p*_1_ and *p*_2_.

### 2.3 Testing β in the mediator model and θ in the outcome model

As in (Zhong et al., 2019), we adopt the widely used SKAT method (Wu et al., 2011) to test *β* in the mediator model to accommodate a potentially large number of correlated SNPs.

Our strategy for testing *θ* in the outcome model consists of four steps: (1) formulation of the likelihood function based on the nature of the outcome random variable *Y*, and (2) Laplace approximation of the likelihood function, and (3) application of the coordinate descent algorithm to estimate parameters by maximizing the approximated likelihood function, and (4) calculation of the likelihood ratio statistic. These four steps allow us to test the mediator effect *θ* in the outcome model.

### 2.4 Likelihood function for the outcome model

To reduce the dimensionality of parameters in the outcome model, we adopted a linear mixed model for continuous outcome in our previously developed SMUT method (Zhong et al., 2019). We consider the following GLMM (McCulloch et al., 2008) when the outcome *Y*_*i*_ follows an exponential family distribution.

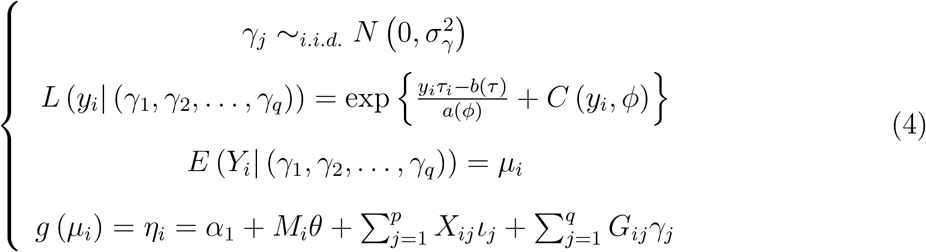

where *τ*_*i*_ is the canonical parameter, *ϕ* the dispersion parameter, *g* the link function, *τ*_*i*_ = *k* (*η*_*i*_) = *b*′^−1^ (*g*^−1^ (*η*_*i*_)).

The likelihood function of the outcome *Y* = (*Y*_1_, *Y*_2_, …, *Y*_*n*_)^*T*^ is

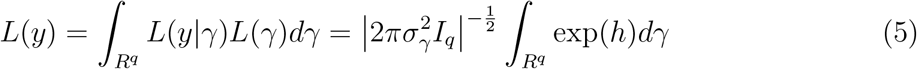

where *L*(*γ*) is the likelihood function for *γ*; 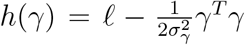 and *ℓ* is the conditional log-likelihood, specifically

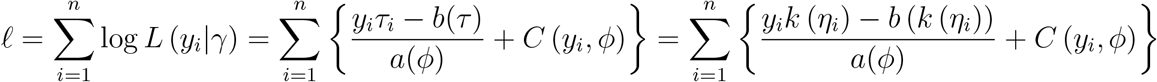

Examples of likelihood function for the outcome from an exponential family distribution are described in the Supplementary Materials Section 1.

When we have a survival outcome, we consider the following mixed effects Cox model (Vaida and Xu, 2000; Pankratz et al., 2005).

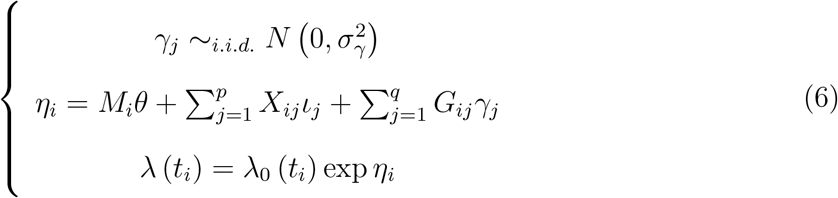

The observed data partial likelihood is

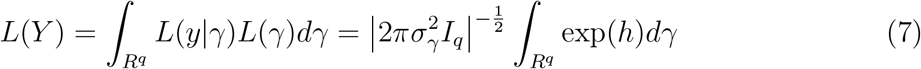

where *L*(*γ*) is the likelihood function for *γ*; 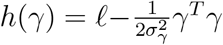; *ℓ* = log *PL* and *PL* is the Cox partial likelihood, specifically 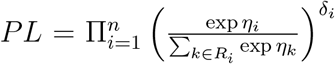 where risk set *R*_*i*_ = {*k* : *Y*_*k*_ ≥ *Y*_*i*_}. Equation 7 takes the same form as equation 5, but the content of the function *h* is different.

### 2.5 Laplace approximation

Laplace’s method is widely applied to approximate the likelihood function (Raudenbush et al., 2000). The integral in equation 5 can be approximated via Laplace’s method by taking Taylor expansion to the second order of *h*(*γ*) around its maximum point 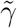.

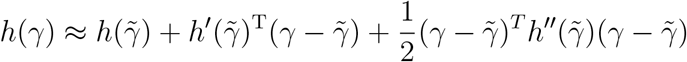

where 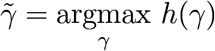. Inserting the Taylor expansion into the integral, we have

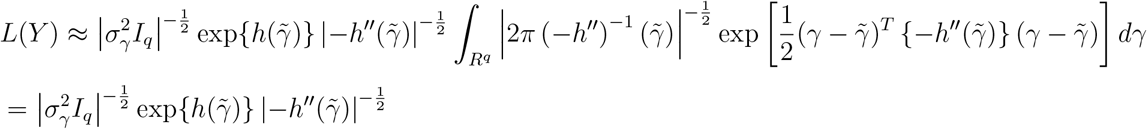

where 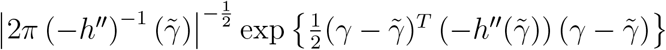 is the probability density function of a multivariate Gaussian distribution, resulting in its integral equal to 1. The approximated log-likelihood *f* is

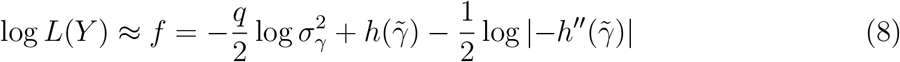

When the outcome *Y* follows an exponential family distribution,

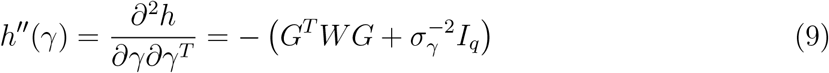

where 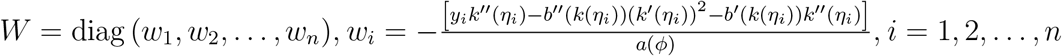.

When we have a survival outcome,

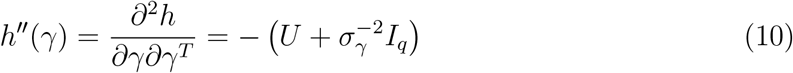

where 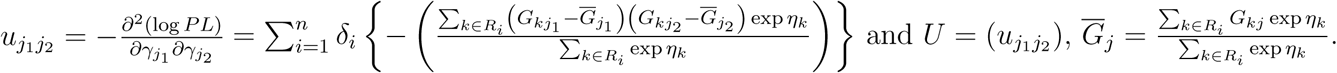

### 2.6 Coordinate descent algorithm

We apply the coordinate descent to maximize the approximated log-likelihood in equation 8. Note that 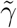 in equation 8 is a function of other parameters, specifically 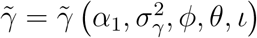. Instead of taking implicit differentiation of 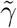 with respect to (w.r.t.) parameters 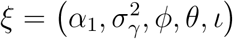 as in (Raudenbush et al., 2000), we use the approximation strategy proposed in (Schelldorfer et al., 2014), which regards 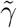 as fixed when updating *ξ*. This strategy is computationally convenient and efficient, at little cost of reduced accuracy. Since 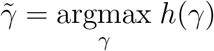, we update 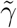 by applying Newton-Raphson algorithm.

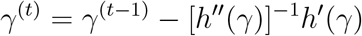

where 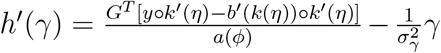, for the outcome following an exponential family distribution; and 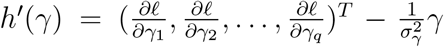 for the survival outcome and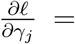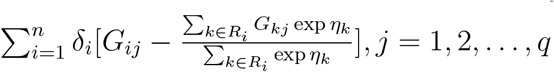.

When taking derivatives of approximated log-likelihood function *f* in equation 8, when the outcome *Y* follows exponential family distribution, we take further approximation by assuming *W* in equation 9 varies slowly as a function of *μ*. This assumption is made in PQL in (Green, 1987; Breslow and Clayton, 1993). When we have a survival outcome, we similarly assume that *U* in equation 10 varies slowly as a function of *η*. Under this assumption, we will only take derivatives of 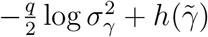 w.r.t. 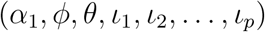 Observe that the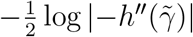 part is only involved when estimating variance component 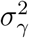. We conduct simulation studies to compare the performance with and without this further approximation.

Assuming 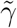 are fixed, we calculate the first and second derivatives of approximated likelihood function *f* as the following.

When the outcome *Y* follows an exponential family distribution, let *ζ* be a vector of (*p* + 2) parameters, 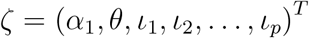. The first derivatives are

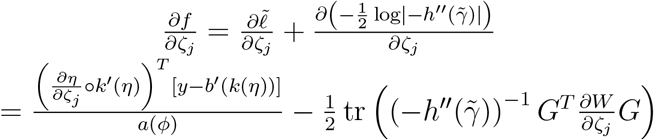

where *j* = 1, 2, …, (*p* + 2) and ◦ is the Hadamard product (entry-wise product), 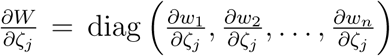 and 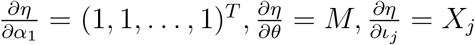.

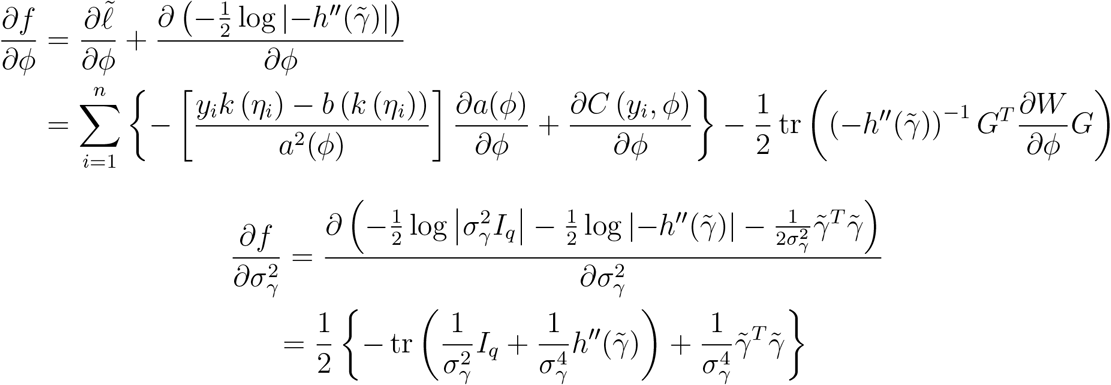

The second derivatives are

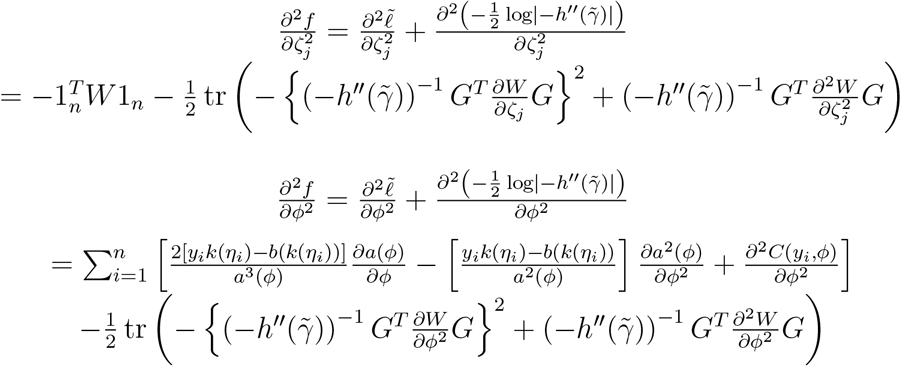

The approximation of derivatives w.r.t. 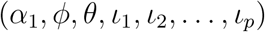 by ignoring the 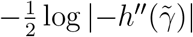 part and assuming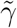 are fixed, is

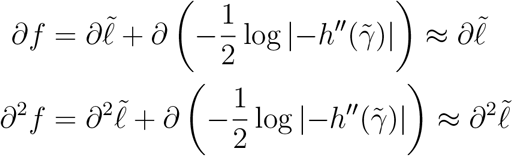

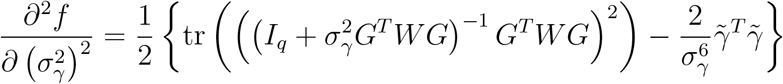

For some commonly used distributions of outcome *Y*, including Gaussian distribution with identity link function, Bernoulli distribution with logit link function, and negative-binomial distribution with log link function, the first and second derivatives of *W* w.r.t. 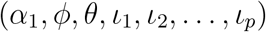 are in the Supplementary Materials Section 2.

When we have a survival outcome, let *ζ* be a vector of (*p*+1) parameters, 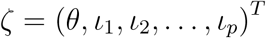

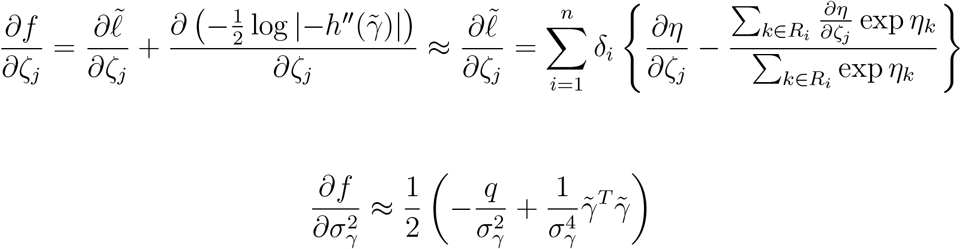

where *j* = 1, 2, …, (*p* + 1) and 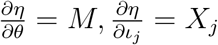.

Because it is computationally intensive to calculate the derivative of 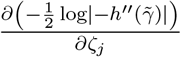, we use 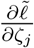 to approximate 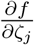.

Finally, we employ the Newton-Raphson algorithm to sequentially update each parameter, say *ψ*, based on their first and second derivatives of *f*.

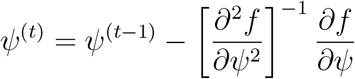

### 2.7 Likelihood ratio test

We obtain approximated likelihood under the null and the alternative hypothesis separately, denoted by *L*_0_ and *L*_1_ respectively. For GLMM, the likelihood ratio statistic 2 (log *L*_1_ − log *L*_0_) asymptotically follows a chi-square distribution with one degree of freedom, and similarly for the partial likelihood ratio statistics for the survival outcome.

## 3. Simulation studies

### 3.1 Simulation settings

To evaluate the performance of SMUT_GLM and SMUT_PH in comparison with alternative methods, we conducted extensive simulations to investigate power and type-I error. Following our previous work (Zhong et al., 2019), we simulated a dataset of 10,000 pseudo-individuals measured at 2,891 SNPs with minor allele frequency (MAF) ≥ 1% in a 1Mb region using the COSI coalescent model (Schaffner et al., 2005) to generate realistic genetic data. The 10,000 pseudo-individuals were constructed by randomly pairing up 20,000 simulated chromosomes without replacement. To evaluate power and type-I error, we generated 500 datasets with 1,000 samples each by sampling without replacement from the entire pool of 10,000 samples simulated above.

The mediator *M* and the outcome *Y* were generated via equations 11. We considered two covariates: one is a continuous variable generated from standard Gaussian distribution and the other is a binary variable generated from Bernoulli(0.5).

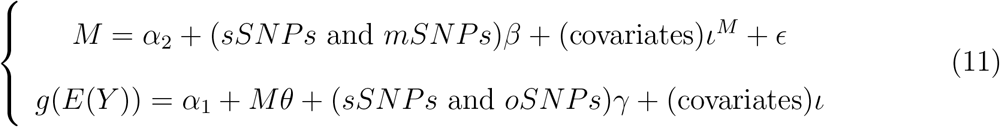

where *g* is the link function and is equal to logit function for binary outcome and log function for count outcome; *ϵ* ∼ *N*(0, 1), *β*_*j*_ ∼_*i.i.d.*_ *c*_*β*_*N* (0, 1), *γ*_*j*_ ∼_*i.i.d.*_ *c*_*γ*_*N* (0, 1), *j* = 1, 2, …, *q*.

We set *c*_*γ*_ = 0.2. The shared SNPs (sSNPs) between the two models are those that influence both the mediator and the outcome. The outcome (or mediator) specific SNPs (oSNPs and mSNPs respectively) only contribute to the outcome (or mediator). The causal SNPs are the union of sSNPs, mSNPs, and oSNPs. We considered two scenarios in terms of causal SNP density: sparse and dense (Table 1). For binary or count outcome, sample size is 1,000 and there are 10 and 500 causal SNPs for sparse and dense scenarios, respectively. For time-to-event outcome, sample size is 200 and there are 10 and 150 causal SNPs for sparse and dense scenarios, respectively. The set of causal SNPs, common across the 500 simulated datasets, were randomly selected from the 2,891 SNPs with MAF ≥ 1%. *β* and *γ*, again fixed across the 500 datasets, were independently drawn from a Gaussian distribution. Error term *ϵ* was independently generated from standard Gaussian distribution and was separately simulated for each of the 500 datasets.

**Table 1.** Causal SNP composition in two simulated scenarios. The sparse(dense) scenario is to simulate data sets based on a small(large) number of causal SNPs. Causal SNPs are the union of shared SNPs, mediator specific SNPs and outcome specific SNPs. Shared SNPs have effects on both mediator and outcome. Mediator(outcome) specific SNPs have effects only on mediator(outcome). All these SNPs are randomly selected from the 2,891 SNPs with MAF ≥ 1%.

| Type of outcome | Sample size | Sparse or dense | # causal SNPs | # sSNPs | # mSNPs | # oSNPs | # non-causal SNPs |
| --- | --- | --- | --- | --- | --- | --- | --- |
| Binary or count | 1000 | Sparse | 10 | 4 | 3 | 3 | 890 |
|  |  | Dense | 500 | 300 | 100 | 100 | 400 |
| Time-to-event | 200 | Sparse | 10 | 4 | 3 | 3 | 190 |
|  |  | Dense | 150 | 90 | 30 | 30 | 50 |

In the simulations, we tested the joint mediation effects of these SNPs on the binary, count or survival outcome using SMUT_GLM and SMUT_PH, as well as other methods including the adapted Huang et al.’s method, adapted LASSO (Tibshirani, 1996). In order to compare the performance of approximations that we adopted, we considered two versions of our method, both treating 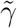 as fixed: (1) based on exact derivatives; (2) based on approximated derivatives. For an outcome from an exponential family distribution, we refer to these two versions as SMUT_GLM exact and SMUT_GLM approxi. For a survival outcome, we refer to the approximated version as SMUT_PH approxi. The exact version of SMUT_PH is not employed because it is hard to derive analytically. The Huang et al.’s method only tests mediator effect in the outcome model, assuming a priori the presence of SNPs’ effects on mediator (i.e., non-zero *β*), adopting a kernel framework where mediator(s) of interest are treated as random and SNPs as fixed (Huang et al., 2015), in contrast to our outcome model where SNPs are treated as random and mediator of interest as fixed. For fair comparison across methods, i.e., testing both *β* and *θ*, we applied the original Huang et al.’s method for the outcome model and SKAT for the mediator model, then combined tests from the two models via IUT, integrating the variance component score test in the outcome model (from the original Huang et al’s method) and score test from SKAT in the mediator model. The adapted LASSO employs LASSO for variable selection in the outcome model and applies IUT using regular regression with the selected variables in the outcome model and all the variables (i.e., genetic variants) via SKAT framework in the mediator model. We applied and compared with the adapted versions of Huang et al. and LASSO because the fnonding original methods only test *θ* in the outcome model. For all the adapted versions, we utilize SKAT to test *β* in the mediator model to be maximally comparable with our SMUT_GLM and SMUT_PH. In other words, adapted Huang et al. is SKAT + original Huang et al. with SKAT fnonding to the testing strategy in the mediator model and original Huang et al. to the testing strategy in the outcome model. Similarly, for LASSO, we use adapted LASSO and SKAT+LASSO exchangeably.

### 3.2 Type-I error in simulations

We evaluated the validity of SMUT_GLM and SMUT_PH along with alternative methods in simulations. SMUT_GLM and SMUT_PH exhibited controlled type-I error rates, at *α* = 0.05 level, regardless of causal SNP density and types of outcome, as shown in Figures 1 and 2 for binary outcome in sparse and dense scenarios respectively, Figures 3 and 4 for time-to-event outcome in sparse and dense scenarios respectively, Supplementary Figures S1 and S2 for count outcome in sparse and dense scenarios respectively. In each figure, the first panel (*c*_*β*_ = 0) and the leftmost point (*θ* = 0) in other panels (*c*_*β*_ ≠ 0) all fnond to the null of no mediation of the SNPs through the mediator. Adapted LASSO and adapted Huang et al.’s method also showed protected type-I error.

**Figure 1.**
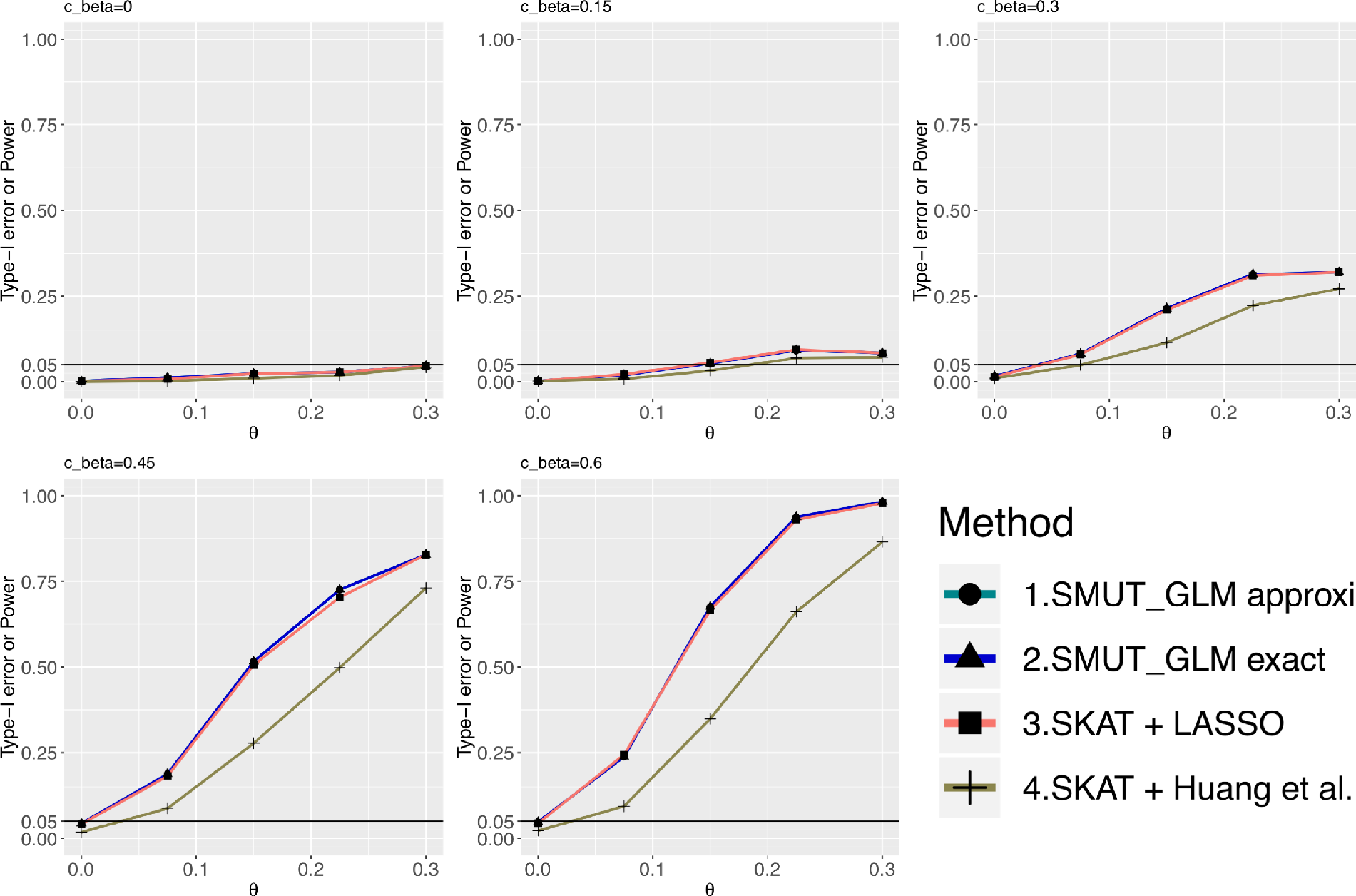
For binary outcome, power and type-I error under sparse causal SNPs scenario. The x-axis is the true mediator effect(*θ*) on the outcome. The y-axis is the power or type-I error. Sub-figures vary in *c*_*β*_ value. *c*_*β*_ = 0 (top-left sub-figure) or *θ* = 0 (left-most points in each sub-figure) are null settings where y-axis represents the fnonding type-I error. When *c*_*β*_ ≠ 0 and *θ* ≠ 0, it is under alternative hypothesis and y-axis represents the fnonding power.

**Figure 2.**
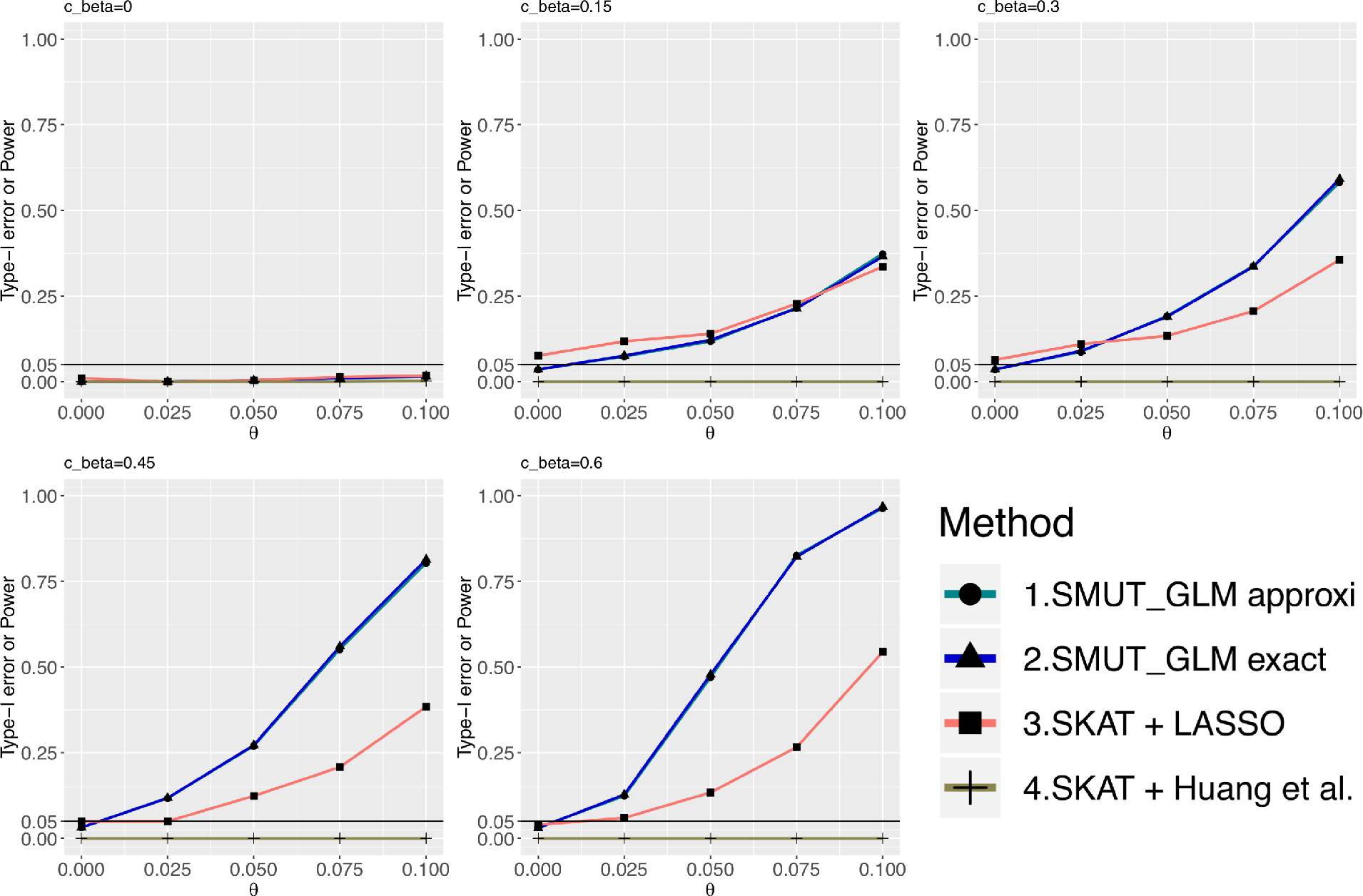
For binary outcome, power and type-I error under dense causal SNPs scenario. X-axis and y-axis are the same as in Figure 1.

**Figure 3.**
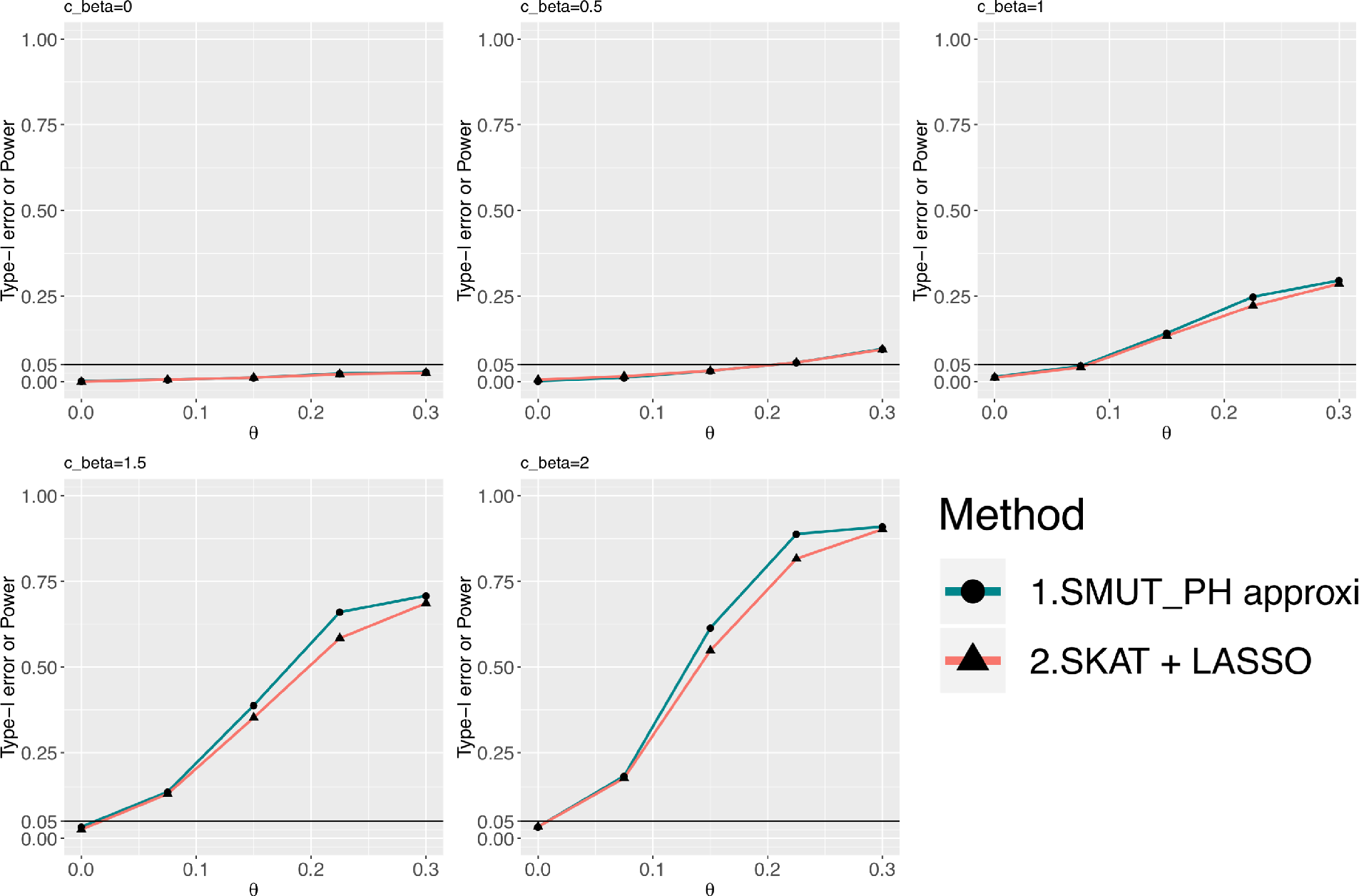
For time-to-event outcome, power and type-I error under sparse causal SNPs scenario. X-axis and y-axis are the same as in Figure 1.

**Figure 4.**
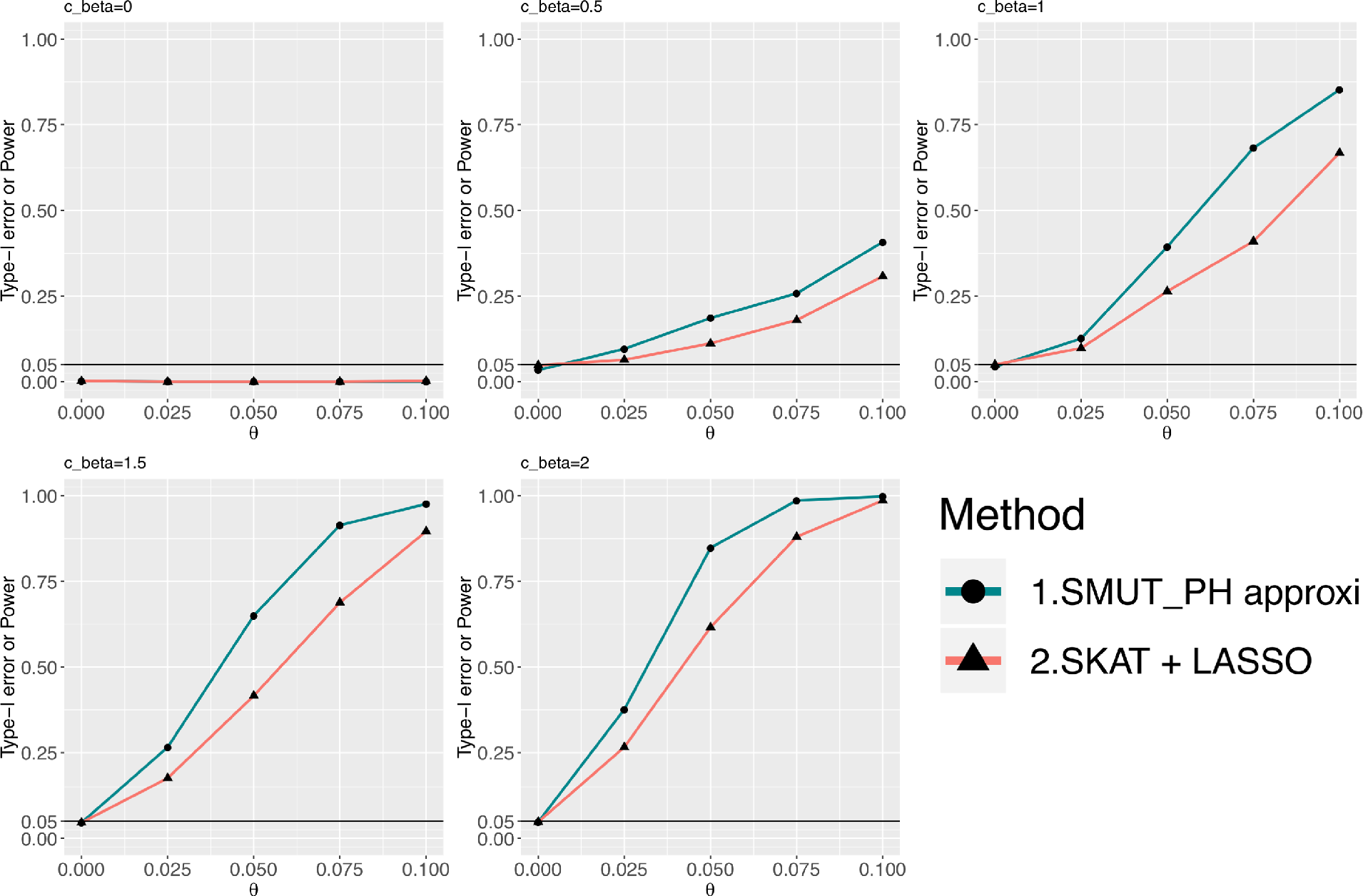
For time-to-event outcome, power and type-I error under dense causal SNPs scenario. X-axis and y-axis are the same as in Figure 1.

### 3.3 Power in simulations

SMUT_GLM and SMUT_PH demonstrated substantial power gains under both the sparse or dense scenarios. We also observe that the approximated version of SMUT_GLM demonstrated very similar performance when compared with its exact counterpart. For example, for a binary outcome and under the scenario of dense causal SNPs when *c*_*β*_ = 0.6, *θ* = 0.1, exact SMUT_GLM, approximated SMUT_GLM, adapted LASSO and adapted Huang et al. had 97%, 96%, 54% and 0% power, respectively. Thus, the power gain, compared with adapted LASSO, was 43% and 42% for exact SMUT_GLM and approximated SMUT_GLM, respectively; and the power gain, compared with adapted Huang et al., was 97% and 96% for exact SMUT_GLM and approximated SMUT_GLM, respectively. For survival outcome, under the scenario of dense causal SNPs when *c*_*β*_ = 1, *θ* = 0.075, approximated SMUT_PH and adapted LASSO had 69% and 41% power, respectively, leading to a power gain of 28%. In addition, power gains appeared more profound with increasing *c*_*β*_ likely because adapted LASSO and adapted Huang et al. becomes more conservative as the pleiotropy effect of SNPs on mediator and outcome (measured by *c*_*β*_) increases.

## 4. Real data application

We assessed our methods and alternatives in real data from two clinical cohorts, which were designed for the study of chlamydia infection. *Chlamydia trachomatis* can ascend from the cervix to the uterus and fallopian tubes in some women, potentially resulting in pelvic inflammatory disease (PID) and severe reproductive morbidities, including infertility and ectopic pregnancy. Recurrent infection leads to worse disease. The first cohort is the T cell Response Against Chlamydia (TRAC) cohort which included asymptomatic women (age 15-30 years) at high risk for sexually transmitted infection (Russell et al., 2015). The second cohort is the Anaerobes and Clearance of Endometritis (ACE) cohort which included symptomatic women (age 15-40 years) with clinically diagnosed PID (Workowski and Bolan, 2015). We analyzed genotype, gene expression and phenotype data of 200 participants combined from these two cohorts. The Institutional Review Boards for Human Subject Research at the University of Pittsburgh and the University of North Carolina approved the study and all participants provided written informed consent prior to inclusion.

### 4.1 Binary outcome

The outcome of interest is ascending chlamydia infection, among participants who had chlamydia infection at enrollment. The control group is the 71 participants who had chlamydia infection restricted to the cervix, and the case group is the 72 participants with both cervical and endometrial chlamydia infection at enrollment. We analyzed genotype, gene expression and phenotype data from these 143 participants.

Here, we tested two genes, *SOS1* and *CD151* gene, for their mediation effects. Son of sevenless homolog 1 (*SOS1*) is a guanine nucleotide exchange factor that in humans is encoded by the *SOS1* gene. The importance of *SOS1* for chlamydia invasion of host cells has been indicated by multiple biomedical studies (Carabeo et al., 2007; Lane et al., 2008; Hackstadt, 2012; Bastidas et al., 2013; Mehlitz and Rudel, 2013; Elwell et al., 2016). The *CD151* gene encodes a protein that is known to complex with integrins. It promotes cell adhesion and may regulate integrin trafficking and/or function. It is a member of the tetraspanin family, which are considered as the gateways for infection (Hauck and Meyer, 2003; Hemler, 2008; Hassuna et al., 2009; Join-Lambert et al., 2010; N Monk and J Partridge, 2012; Seu et al., 2017). In addition, SNPs annotation database, RegulomeDB (Boyle et al., 2012), demonstrates that some SNPs in these two genes are eQTLs with experimental evidence. Thus, the presence of mediation effect via the expression of each gene is expected.

We first extracted SNPs within ± 1 Mb of the fnonding genes and then conducted expression quantitative trait loci (eQTLs) analysis for these two genes. The eQTL analysis was conducted based on all the 200 participants. For the first gene *SOS1*, mediation testing encompassed 83 SNPs with MAF ≥ 10% and significant eQTL association (with *SOS1*) at a false discovery rate (FDR) threshold of 10%, using SMUT_GLM, adapted LASSO and adapted Huang et al.’s method. Both SMUT_GLM and adapted Huang et al.’s method detected significant mediation effects, while adapted LASSO did not (Table 2). For the second gene *CD151*, our mediation (via expression of *CD151*) testing involved 40 SNPs with MAF ≥ 10% and significant eQTL (with *CD151*) at FDR 10%. Only SMUT_GLM showed significant mediation effects of these SNPs through the expression of *CD151* on ascending chlamydia infection (Table 2).

**Table 2.** Real data application to TRAC and ACE datasets.

|  |  |  |  |  | *P values* |  |
| --- | --- | --- | --- | --- | --- | --- |
| Type of outcome | Gene | Probesets | #SNPs | SMUT_GLM | LASSO | Huang et al. |
| Binary | *SOS1* | 2140519 | 83 | 0.0235 | 0.0691 | 0.0229 |
| Binary | *CD151* | 1940132 | 40 | 0.0245 | 0.1192 | 0.2289 |
|  |  |  |  | SMUT_PH | LASSO | Huang et al. |
| Time-to-event | *BIRC3* | 7210154 | 4 | 0.001 | 0.001 | 0.002 |

### 4.2 Time-to-event outcome

TRAC participants returned for follow-up visits at 1, 4, 8, and 12 months after enrollment. The outcome of interest we evaluated here is time to the first incident chlamydia infection. We analyzed genotype, gene expression and time-to-event data from all 181 participants in the TRAC cohort who had both genotype and gene expression data available.

Here, we tested a gene, *BIRC3*, for its mediation effect. The gene *BIRC3* encodes for Baculoviral IAP Repeat Containing 3, a E3 ubiquitin-protein ligase regulating NF-kappa-B signaling (Blankenship et al., 2009; Kim et al., 2010; Tan et al., 2013). It acts as an important regulator of pathogen recognition receptor signaling (Bertrand et al., 2009), which can have profound effects on the development of downstream adaptive immune responses (Takeda et al., 2003; Palm and Medzhitov, 2009; Kumar et al., 2011). In addition, biological studies suggested that *BIRC3* may protect mammalian host cells against apoptosis, leading to accommodate chlamydial growth (Bryant et al., 2004; Park et al., 2004; Paland et al., 2006; Ying et al., 2008). Therefore, mediation effect via the expression of BIRC3 gene is logical. Our mediation testing involved 4 SNPs with MAF ≥ 10% and eQTL (with *BIRC3* at FDR 10%, using SMUT_PH, adapted LASSO and adapted Huang et al.’s method. All the methods showed significant mediation effects through *BIRC3* on incident chlamydia infection (Table 2).

## 5. Discussion

Our proposed methods, SMUT_GLM and SMUT_PH, extend our previous work (Zhong et al., 2019) to test mediation effect of multiple correlated genetic variants on a non-Gaussian outcome (e.g. binary, count, time-to-event outcome) through a mediator (e.g. expression of some gene in the vicinity). We employ intersection-union test approach to derive a single *p* value by integrating *p* values from separate tests of *β* and *θ*. Moreover, our methods do not rely on complete mediation assumption nor presume independent genetic variants. SMUT_GLM and SMUT_PH are statistically more powerful than alternative methods including adapted LASSO and adapted Huang et al.’s method. Power loss using adapted LASSO might be a result of violating its sparsity assumption. Huang et al.’s method has lower statistical power, which might be due to the modeling strategy for the mediator effect (*θ*). Specifically, the mediator effect (*θ*) is modeled as a random effect in the outcome model by Huang et al.’s method, which might not be optimal particularly when only one mediator is considered at a time. When jointly testing multiple mediators in the outcome model, Huang et al.’s method may perform more favorably. However, testing one mediator at a time has the advantage to pinpoint or prioritize the causal gene(s), which would not be possible when testing multiple genes in aggregate via Huang et al.’s random mediator effect.

One limitation of our proposed methods is that we assume the effects of genetic variants follow a Gaussian distribution. This may not be correct when there are non-causal SNPs in the model and in this case, a mixture distribution might be more appropriate. It is reassuring to observe protected type-I error from our simulation studies, which included considerable number of proportion of non-causal SNPs in all scenarios considered. More properly modeling the effects of genetic variants may further increase the statistical power under the alternative hypotheses but due to modeling complexity and subsequently inevitable computational costs, we decide not to further pursue this in our current work. This is an interesting topic for future investigation.

Our proposed methods can be further extended to handle multiple correlated outcomes to gain additional power. One possible approach to model correlation among multiple outcomes is to add random intercepts in the outcome model. When adding random intercepts to the model, additional Laplace approximation will be applied to these random effects. The accuracy of Laplace approximation by taking only the second order of the Taylor expansion needs further investigation in such more complicated model. If second order is insufficient, higher-order of Laplace approximation (Raudenbush et al., 2000) could be considered to achieve higher precision, at the cost of increased computational burden, which can be high with high-dimensional random effects. Such work thus warrants separate investigation and a separate publication.

In simulation studies, we also compared the computational time of our methods with adapted LASSO and adapted Huang et al.’s method. In general, our methods’ computational time is similar to that of adapted LASSO, and both our methods and adapted LASSO run faster than Huang et al.’s method (Supplementary Figure S3, S4, S5). Our methods use approximations when calculating derivatives of the likelihood functions, which substantially reduces computational burden (Supplementary Figure S3, S4). For the binary and count outcome with sparse causal SNPs, our SMUT_GLM runs faster than adapted LASSO. For the binary and count outcome with dense causal SNPs, our method runs more slowly than adapted LASSO. We suspect that our method takes longer time to converge under the dense scenario than under the sparse scenario because there are more non-zero coefficients under the dense scenario.

In summary, we proposed SMUT_GLM and SMUT_PH that can test mediation effects of multiple correlated genetic variants on a non-Gaussian outcome through a mediator. We anticipate our proposed method will become a powerful tool to bridge the gap in terms of molecular mechanisms between various types of phenotypes and the fnonding associated genetic variant(s) identified in recent literature.

## ACKNOWLEDGEMENTS

This work was supported by the National Institutes of Health U19 AI084024 and R01 AI119164 to TD. We thank all participants in ACE and TRAC for agreeing to take part in the studies, and all investigators in these two studies for sharing the data.

## SUPPLEMENTARY MATERIALS

The Supplementary Figures and Results referenced in Sections 2, 3, and 5 are available with this article online. The R code for our method is the function GSMUT in the R package SMUT, which is publicly available from CRAN at https://CRAN.R-project.org/package=SMUT.

## REFERENCES

Baron, R. M. and Kenny, D. a. (1986). The moderator-mediator variable distinction in social psychological research: conceptual, strategic, and statistical considerations. Journal of personality and social psychology 51, 1173–1182.

Bastidas, R. J., Elwell, C. A., Engel, J. N., and Valdivia, R. H. (2013). Chlamydial intracellular survival strategies. Cold Spring Harbor perspectives in medicine 3, a010256.

Berger, R. L. and Hsu, J. C. (1996). Bioequivalence trials, intersection-union tests and equivalence confidence sets. Statistical Science 11, 283–319.

Bertrand, M. J., Doiron, K., Labbé, K., Korneluk, R. G., Barker, P. A., and Saleh, M. (2009). Cellular inhibitors of apoptosis ciap1 and ciap2 are required for innate immunity signaling by the pattern recognition receptors nod1 and nod2. Immunity 30, 789–801.

Blankenship, J. W., Varfolomeev, E., Goncharov, T., Fedorova, A. V., Kirkpatrick, D. S., Izrael-Tomasevic, A., Phu, L., Arnott, D., Aghajan, M., Zobel, K., et al. (2009). Ubiquitin binding modulates iap antagonist-stimulated proteasomal degradation of c-iap1 and c-iap2 1. Biochemical Journal 417, 149–165.

Bolker, B. M., Brooks, M. E., Clark, C. J., Geange, S. W., Poulsen, J. R., Stevens, M. H. H., and White, J.-S. S. (2009). Generalized linear mixed models: a practical guide for ecology and evolution. Trends in ecology & evolution 24, 127–135.

Boyle, A. P., Hong, E. L., Hariharan, M., Cheng, Y., Schaub, M. A., Kasowski, M., Karczewski, K. J., Park, J., Hitz, B. C., Weng, S., et al. (2012). Annotation of functional variation in personal genomes using regulomedb. Genome research 22, 1790–1797.

Breslow, N. E. and Clayton, D. G. (1993). Approximate Inference in Generalized Linear Mixed Models. Journal of the American Statistical Association 88, 9.

Bryant, P. A., Venter, D., Robins-Browne, R., and Curtis, N. (2004). Chips with everything: Dna microarrays in infectious diseases. The Lancet infectious diseases 4, 100–111.

Carabeo, R. A., Dooley, C. A., Grieshaber, S. S., and Hackstadt, T. (2007). Rac interacts with abi-1 and wave2 to promote an arp2/3-dependent actin recruitment during chlamydial invasion. Cellular microbiology 9, 2278–2288.

Chen, H., Huffman, J. E., Brody, J. A., Wang, C., Lee, S., Li, Z., Gogarten, S. M., Sofer, T., Bielak, L. F., Bis, J. C., et al. (2019). Efficient variant set mixed model association tests for continuous and binary traits in large-scale whole-genome sequencing studies. The American Journal of Human Genetics 104, 260–274.

Chen, H., Wang, C., Conomos, M. P., Stilp, A. M., Li, Z., Sofer, T., Szpiro, A. A., Chen, W., Brehm, J. M., Celedón, J. C., et al. (2016). Control for population structure and relatedness for binary traits in genetic association studies via logistic mixed models. The American Journal of Human Genetics 98, 653–666.

Cheng, J., Cheng, N. F., Guo, Z., Gregorich, S., Ismail, A. I., and Gansky, S. A. (2018). Mediation analysis for count and zero-inflated count data. Statistical methods in medical research 27, 2756–2774.

Daubechies, I., Defrise, M., and De Mol, C. (2004). An iterative thresholding algorithm for linear inverse problems with a sparsity constraint. Communications on Pure and Applied Mathematics: A Journal Issued by the Courant Institute of Mathematical Sciences 57, 1413–1457.

Elwell, C., Mirrashidi, K., and Engel, J. (2016). Chlamydia cell biology and pathogenesis. Nature Reviews Microbiology 14, 385.

Friedman, J., Hastie, T., and Tibshirani, R. (2008). Sparse inverse covariance estimation with the graphical lasso. Biostatistics 9, 432–441.

Friedman, J., Hastie, T., and Tibshirani, R. (2010). Regularization paths for generalized linear models via coordinate descent. Journal of statistical software 33, 1.

Fu, W. J. (1998). Penalized regressions: the bridge versus the lasso. Journal of computational and graphical statistics 7, 397–416.

Gilks, W. R. (1996). Introducing markov chain monte carlo. Markov Chain Monte Carlo in Practice.

Green, P. J. (1987). Penalized Likelihood for General Semi-Parametric Regression Models. International Statistical Review 55, 245–259.

Hackstadt, T. (2012). Intracellular pathogens I: chlamydiales, volume 1, pages 126–148. American Society for Microbiology Press.

Hassuna, N., Monk, P. N., Moseley, G. W., and Partridge, L. J. (2009). Strategies for targeting tetraspanin proteins. BioDrugs 23, 341–359.

Hauck, C. R. and Meyer, T. F. (2003). smalltalk: Opa proteins as mediators of neisseria– host-cell communication. Current opinion in microbiology 6, 43–49.

Hemler, M. E. (2008). Targeting of tetraspanin proteinspotential benefits and strategies. Nature reviews Drug discovery 7, 747.

Huang, Y.-T., Cai, T., and Kim, E. (2016). Integrative genomic testing of cancer survival using semiparametric linear transformation models. Statistics in medicine 35, 2831–44.

Huang, Y.-T., Liang, L., Moffatt, M. F., Cookson, W. O. C. M., and Lin, X. (2015). iGWAS: Integrative Genome-Wide Association Studies of Genetic and Genomic Data for Disease Susceptibility Using Mediation Analysis. Genetic Epidemiology 39, 347–356.

Huang, Y.-T. and Pan, W.-C. (2016). Hypothesis test of mediation effect in causal mediation model with high-dimensional continuous mediators. Biometrics 72, 402–413.

Join-Lambert, O., Morand, P. C., Carbonnelle, E., Coureuil, M., Bille, E., Bourdoulous, S., and Nassif, X. (2010). Mechanisms of meningeal invasion by a bacterial extracellular pathogen, the example of neisseria meningitidis. Progress in neurobiology 91, 130–139.

Kim, C. W., Kim, H. K., Vo, M.-T., Lee, H. H., Kim, H. J., Min, Y. J., Cho, W. J., and Park, J. W. (2010). Tristetraprolin controls the stability of ciap2 mrna through binding to the 3 utr of ciap2 mrna. Biochemical and biophysical research communications 400, 46–52.

Kumar, H., Kawai, T., and Akira, S. (2011). Pathogen recognition by the innate immune system. International reviews of immunology 30, 16–34.

Lane, B. J., Mutchler, C., Al Khodor, S., Grieshaber, S. S., and Carabeo, R. A. (2008). Chlamydial entry involves tarp binding of guanine nucleotide exchange factors. PLoS pathogens 4, e1000014.

MacKinnon, D. P., Fairchild, A. J., and Fritz, M. S. (2007). Mediation Analysis. Annual Review of Psychology 58, 593–614.

McCullagh, P. and Nelder, J. (1989). Generalized Linear Models. CRC Press Vol. 37 pages 1–514.

McCulloch, C. E. and Searle, S. R. (2001). Generalized, Linear, and Mixed Models, volume 45.

McCulloch, C. E., Searle, S. R., and Neuhaus, J. M. (2008). Generalized, Linear, and Mixed Models, 2nd Edition.

Mehlitz, A. and Rudel, T. (2013). Modulation of host signaling and cellular responses by chlamydia. Cell Communication and Signaling 11, 90.

N Monk, P. and J Partridge, L. (2012). Tetraspanins-gateways for infection. Infectious Disorders-Drug Targets (Formerly Current Drug Targets-Infectious Disorders) 12, 4–17.

O’Rourke, H. P. and Vazquez, E. (2019). Mediation analysis with zero-inflated substance use outcomes: Challenges and recommendations. Addictive behaviors.

Paland, N., Rajalingam, K., Machuy, N., Szczepek, A., Wehrl, W., and Rudel, T. (2006). Nf-*κ*b and inhibitor of apoptosis proteins are required for apoptosis resistance of epithelial cells persistently infected with chlamydophila pneumoniae. Cellular microbiology 8, 1643–1655.

Palm, N. W. and Medzhitov, R. (2009). Pattern recognition receptors and control of adaptive immunity. Immunological reviews 227, 221–233.

Pankratz, V. S., De Andrade, M., and Therneau, T. M. (2005). Random-effects cox proportional hazards model: General variance components methods for time-to-event data. Genetic Epidemiology: The Official Publication of the International Genetic Epidemiology Society 28, 97–109.

Park, J. Y., Wu, C., Basu, S., McGue, M., and Pan, W. (2018). Adaptive snp-set association testing in generalized linear mixed models with application to family studies. Behavior genetics 48, 55–66.

Park, S.-M., Yoon, J.-B., and Lee, T. H. (2004). Receptor interacting protein is ubiquitinated by cellular inhibitor of apoptosis proteins (c-iap1 and c-iap2) in vitro. FEBS letters 566, 151–156.

Preacher, K. J. (2015). Advances in mediation analysis: A survey and synthesis of new developments. Annual review of psychology 66, 825–852.

Raudenbush, S. W., Yang, M. L., and Yosef, M. (2000). Maximum Likelihood for Generalized Linear Models with Nested Random Effects via High-Order, Multivariate Laplace Approximation. Journal of Computational and Graphical Statistics 9, 141–157.

Russell, A. N., Zheng, X., O’connell, C. M., Taylor, B. D., Wiesenfeld, H. C., Hillier, S. L., Zhong, W., and Darville, T. (2015). Analysis of factors driving incident and ascending infection and the role of serum antibody in Chlamydia trachomatis genital tract infection. The Journal of infectious diseases 213, 523–531.

Schaffner, S. F., Foo, C., Gabriel, S., Reich, D., Daly, M. J., and Altshuler, D. (2005). Calibrating a coalescent simulation of human genome sequence variation. Genome Research 15, 1576–1583.

Schelldorfer, J., Meier, L., and Bühlmann, P. (2014). Glmmlasso: an algorithm for high-dimensional generalized linear mixed models using 1-penalization. Journal of Computational and Graphical Statistics 23, 460–477.

Seu, L., Tidwell, C., Timares, L., Duverger, A., Wagner, F. H., Goepfert, P. A., Westfall, A. O., Sabbaj, S., and Kutsch, O. (2017). Cd151 expression is associated with a hyperproliferative t cell phenotype. The Journal of Immunology 199, 3336–3347.

Takeda, K., Kaisho, T., and Akira, S. (2003). Toll-like receptors. Annual review of immunology 21, 335–376.

Tan, B., Zammit, N. W., Yam, A. O., Slattery, R., Walters, S., Malle, E., and Grey, S. (2013). Baculoviral inhibitors of apoptosis repeat containing (birc) proteins fine-tune tnf-induced nuclear factor *κ*b and c-jun n-terminal kinase signalling in mouse pancreatic beta cells. Diabetologia 56, 520–532.

Tibshirani, R. (1996). Regression shrinkage and selection via the lasso. Journal of the Royal Statistical Society. Series B (Methodological) pages 267–288.

Vaida, F. and Xu, R. (2000). Proportional hazards model with random effects. Statistics in medicine 19, 3309–3324.

Workowski, K. A. and Bolan, G. A. (2015). Sexually transmitted diseases treatment guidelines, 2015. MMWR. Recommendations and reports: Morbidity and mortality weekly report. Recommendations and reports 64, 1.

Wu, M. C., Lee, S., Cai, T., Li, Y., Boehnke, M., and Lin, X. (2011). Rare-variant association testing for sequencing data with the sequence kernel association test. American Journal of Human Genetics 89, 82–93.

Yan, Q., Tiwari, H. K., Yi, N., Gao, G., Zhang, K., Lin, W.-Y., Lou, X.-Y., Cui, X., and Liu, N. (2015). A sequence kernel association test for dichotomous traits in family samples under a generalized linear mixed model. Human heredity 79, 60–68.

Ying, S., Christian, J. G., Paschen, S. A., and Häcker, G. (2008). Chlamydia trachomatis can protect host cells against apoptosis in the absence of cellular inhibitor of apoptosis proteins and mcl-1. Microbes and infection 10, 97–101.

Zhang, H., Zheng, Y., Zhang, Z., Gao, T., Joyce, B., Yoon, G., Zhang, W., Schwartz, J., Just, A., Colicino, E., et al. (2016). Estimating and testing high-dimensional mediation effects in epigenetic studies. Bioinformatics 32, 3150–3154.

Zhong, W., Spracklen, C. N., Mohlke, K. L., Zheng, X., Fine, J., and Li, Y. (2019). Multi-snp mediation intersection-union test. Bioinformatics.

