## Supplementary Materials for "Generalized Multi-SNP Mediation Intersection-Union Test"

### Section 1 Likelihood function for the outcome from an exponential family distribution.

**For continuous outcome $Y$ (Gaussian distribution with identity link function)**

$$L(y_i|\gamma) = \exp\left(\frac{y_i\mu_i - \frac{1}{2}{\mu_i}^2}{\sigma^2} - \frac{y_i^2}{2\sigma^2} - \frac{1}{2}\log\sigma^2\right)$$

where the constant term is ignored in the likelihood function.

$$g\,(x) = x, \eta_i = \mu_i = \tau_i, \phi\ = \sigma^2, a\,(x) = x$$

$$C\ = -\frac{Y_i^2}{2\phi} - \frac{1}{2}\log\phi$$

$$b\,(\tau_i) = \frac{\mu_i^2}{2}\,, b\,(x) = \frac{x^2}{2}, b'\,(x) = x\,, b''\,(x) = 1$$

$$k\,(x) = b\,'^{-1}\big(g^{\,-1}(x)\big) = b\,'^{-1}(x) = x$$

$$k\,'(x) = 1\,, k\,''(x) = 0$$

weight $w_i = \frac{1}{\phi}\,, W\ = diag(w_1, w_2, \dots, w_n)$

**For binary outcome $Y$ (Bernoulli distribution with logit link function)**

$$L(y_i|\gamma) = \exp\left(\frac{y_i\log\left(\frac{\mu_i}{1-\mu_i}\right) + \log(1-\mu_i)}{1}\right)$$

$$\tau_i = \log\left(\frac{\mu_i}{1-\mu_i}\right)$$

$$b(\tau_i) = -\log(1-\mu_i) = \log(1+\exp\tau_i)$$

$$a(\phi) = 1\,, C(y_i,\phi) = 0$$

$$E(y_i) = \mu_i$$

$g$ is the canonical link function,

$$g(\mu_i) = logit(\mu_i) = \eta_i = \alpha + M_i\theta + G_{i1}\gamma_1 + \cdots + G_{iq}\gamma_q + X_{i1}\iota_1 + X_{i2}\iota_2 + \cdots + X_{ip}\iota_p$$

$$\gamma_j \sim i.i.d.\ N(0,\sigma_\gamma^2)$$

$$\tau_i = k(\eta_i) = b'^{-1}\big(g^{-1}(\eta_i)\big) = \eta_i$$
$$W = diag(w_1, w_2, \dots, w_n), \text{ and } w_i = b''(\eta_i) = \frac{\exp \eta_i}{(1+\exp \eta_i)^2} = \mu_i(1-\mu_i)$$

**For count outcome $Y$ (Negative-binomial distribution with log link function)**

$$L(y_i|\gamma) = \frac{\Gamma(y_i+\phi)}{\Gamma(\phi)y_i!}\left(\frac{\phi}{\phi+\mu_i}\right)^{\phi}\left(\frac{\mu_i}{\phi+\mu_i}\right)^{y_i}$$
$$= \exp\left[y_i \log\frac{\mu_i}{\phi+\mu_i} + \phi\log\frac{\phi}{\phi+\mu_i} + \log\frac{\Gamma(y_i+\phi)}{\Gamma(\phi)y_i!}\right]$$

Let $\pi_i \coloneqq \frac{\phi}{\phi+\mu_i}$

$$L(y_i|\gamma) = \exp\left[y_i\log(1-\pi_i) + \phi\log\pi_i + \log\frac{\Gamma(y_i+\phi)}{\Gamma(\phi)y_i!}\right]$$
$$\tau_i = \log(1-\pi_i)$$
$$b(\tau_i) = -\phi\log\pi_i = -\phi\log(1-\exp\tau_i)$$
$$a(\phi) = 1, \qquad C(y_i,\phi) = \log\frac{\Gamma(y_i+\phi)}{\Gamma(\phi)y_i!}$$
$$E(y_i) = b'(\tau_i) = \frac{\phi\exp\tau_i}{1-\exp\tau_i} = \mu_i$$
$$b''(\tau_i) = \frac{\phi\exp\tau_i}{(1-\exp\tau_i)^2} = \frac{\mu_i}{\pi_i}$$
$$g(\mu_i) = \log(\mu_i) = \eta_i = \alpha + M_i\theta + G_{i1}\gamma_1 + \cdots + G_{iq}\gamma_q + X_{i1}\iota_1 + X_{i2}\iota_2 + \cdots + X_{ip}\iota_p$$
$$\gamma_j \sim i.i.d.\ N(0,\sigma_\gamma^2)$$
$$\tau_i = k(\eta_i) = b'^{-1}\big(g^{-1}(\eta_i)\big) = \eta_i - \log(\phi+\exp\eta_i)$$
$$\exp\tau_i = \frac{\exp\eta_i}{\phi+\exp\eta_i}$$
$$k'(\eta_i) = \frac{\phi}{\phi+\exp\eta_i} = \pi_i$$
$$k''(\eta_i) = -\frac{\phi\exp\eta_i}{(\phi+\exp\eta_i)^2} = -\pi_i(1-\pi_i)$$

$W = diag(w_1, w_2, \dots, w_n)$, and

$$w_i = -\frac{\left[y_i k''(\eta_i) - b''\big(k(\eta_i)\big)\big(k'(\eta_i)\big)^2 - b'\big(k(\eta_i)\big)k''(\eta_i)\right]}{\phi}$$
$$= -\left[-y_i\pi_i(1-\pi_i) - \frac{\mu_i}{\pi_i}\pi_i^2 + \mu_i\pi_i(1-\pi_i)\right] = y_i\pi_i(1-\pi_i) + \mu_i\pi_i^2$$

**Section 2 Derivatives of $W = diag(w_1, w_2, \dots, w_n)$ w.r.t. $(\alpha_1, \phi, \theta, \iota_1, \iota_2, \dots, \iota_p, \gamma_1, \gamma_2, \dots, \gamma_q)$**

$\eta_i = \alpha_1 + M_i\theta + \Sigma_{j=1}^{p} X_{ij}\tau_j + \Sigma_{j=1}^{q} G_{ij}\iota_j$

Supplementary Table 1. The first derivative of $\eta_i$ w.r.t. parameters

$\xi = (\xi_1, \xi_2, \dots, \xi_{p+q+3})^T = (\alpha_1, \phi, \theta, \iota_1, \iota_2, \dots, \iota_p, \gamma_1, \gamma_2, \dots, \gamma_q)^T$

| Parameter | First derivative of $\eta_i$ |
|---|---|
| $\alpha_1$ | 1 |
| $\theta$ | $M_i$ |
| $\iota_j$ | $X_{ij}$ |
| $\gamma_j$ | $G_{ij}$ |

For continuous outcome (Gaussian distribution with identity link function)

$$w_i = \frac{1}{\phi}, i = 1,2, \dots, n$$

$$\frac{\partial w_i}{\partial \xi_j} = 0, \frac{\partial^2 w_i}{\partial \xi_j^2} = 0$$

$$\frac{\partial w_i}{\partial \phi} = -\frac{1}{\phi^2}$$

$$\frac{\partial^2 w_i}{\partial \phi^2} = \frac{2}{\phi^3}$$

For binary outcome (Bernoulli distribution with logit link function)

$$w_i = \frac{\exp \eta_i}{(1 + \exp \eta_i)^2} = \mu_i(1 - \mu_i)$$

where $\mu_i = \frac{\exp \eta_i}{1+\exp \eta_i}$

$$\frac{\partial w_i}{\partial \xi_j} = \frac{(\exp \eta_i)(1 - \exp \eta_i)}{(1 + \exp \eta_i)^3} \frac{\partial \eta_i}{\partial \xi_j} = w_i(1 - 2\mu_i) \frac{\partial \eta_i}{\partial \xi_j}$$

$\frac{\partial^2 w_i}{\partial {\xi_j}^2} = \exp \eta_i \frac{(1-4\exp \eta_i + (\exp \eta_i)^2)}{(1+\exp \eta_i)^4} \left(\frac{\partial \eta_i}{\partial \xi_j}\right)^2 = w_i(1 - 6w_i) \left(\frac{\partial \eta_i}{\partial \xi_j}\right)^2$, since $\frac{\partial^2 \eta_i}{\partial {\xi_j}^2} = 0$

For count outcome (Negative-binomial distribution with log link function)

$$w_i = y_i\pi_i(1 - \pi_i) + \mu_i\pi_i^2 = y_i \frac{\phi}{\phi + \mu_i} \frac{\mu_i}{\phi + \mu_i} + \mu_i \left(\frac{\phi}{\phi + \mu_i}\right)^2 = (y_i + \phi) \frac{\phi\mu_i}{(\phi + \mu_i)^2}$$
$$= (y_i + \phi)\pi_i(1 - \pi_i)$$

where $\pi_i = \frac{\phi}{\phi + \exp \eta_i}$

$$\frac{\partial w_i}{\partial \xi_j} = (Y_i + \phi) \frac{\partial(\pi_i(1 - \pi_i))}{\partial \xi_j}$$

$$= (Y_i + \phi) \frac{\partial \left( \frac{\phi \exp \eta_i}{(\phi + \exp \eta_i)^2} \right)}{\partial \xi_j}$$

$$= (Y_i + \phi) \frac{(\phi \mu_i)(\phi - \mu_i)}{(\phi + \mu_i)^3} \frac{\partial \eta_i}{\partial \xi_j}$$

$\frac{\partial^2 w_i}{\partial {\xi_j}^2} = (Y_i + \phi)(\phi \mu_i) \frac{(\phi^2 - 4\phi\mu_i + \mu_i^2)}{(\phi + \mu_i)^4} \left( \frac{\partial \eta_i}{\partial \xi_j} \right)^2$, since $\frac{\partial^2 \eta_i}{\partial {\xi_j}^2} = 0$

$$\frac{\partial w_i}{\partial \phi} = \pi_i(1 - \pi_i) + (Y_i + \phi) \frac{\partial(\pi_i(1 - \pi_i))}{\partial \phi}$$

$$= \pi_i(1 - \pi_i) + (Y_i + \phi) \frac{\partial \left( \frac{\phi \exp \eta_i}{(\phi + \exp \eta_i)^2} \right)}{\partial \phi}$$

$$= \pi_i(1 - \pi_i) + (Y_i + \phi) \frac{(\exp \eta_i)(\phi + \exp \eta_i)^2 - 2(\phi \exp \eta_i)(\phi + \exp \eta_i)}{(\phi + \exp \eta_i)^4}$$

$$= \pi_i(1 - \pi_i) + (Y_i + \phi) \frac{(\exp \eta_i)^2 - \phi \exp \eta_i}{(\phi + \exp \eta_i)^3}$$

$$\frac{\partial^2 w_i}{\partial \phi^2} = \frac{\partial(\pi_i(1 - \pi_i))}{\partial \phi} + \frac{(\exp \eta_i)^2 - \phi \exp \eta_i}{(\phi + \exp \eta_i)^3}$$

$$+ (Y_i + \phi) \frac{-(\exp \eta_i)(\phi + \exp \eta_i)^3 - \{(\exp \eta_i)^2 - \phi \exp \eta_i\} 3(\phi + \exp \eta_i)^2}{(\phi + \exp \eta_i)^6}$$

$$= 2 \frac{(\exp \eta_i)^2 - \phi \exp \eta_i}{(\phi + \exp \eta_i)^3} + (Y_i + \phi) \frac{2\phi \exp \eta_i - 4(\exp \eta_i)^2}{(\phi + \exp \eta_i)^4}$$

### Section 3 Supplementary Figures

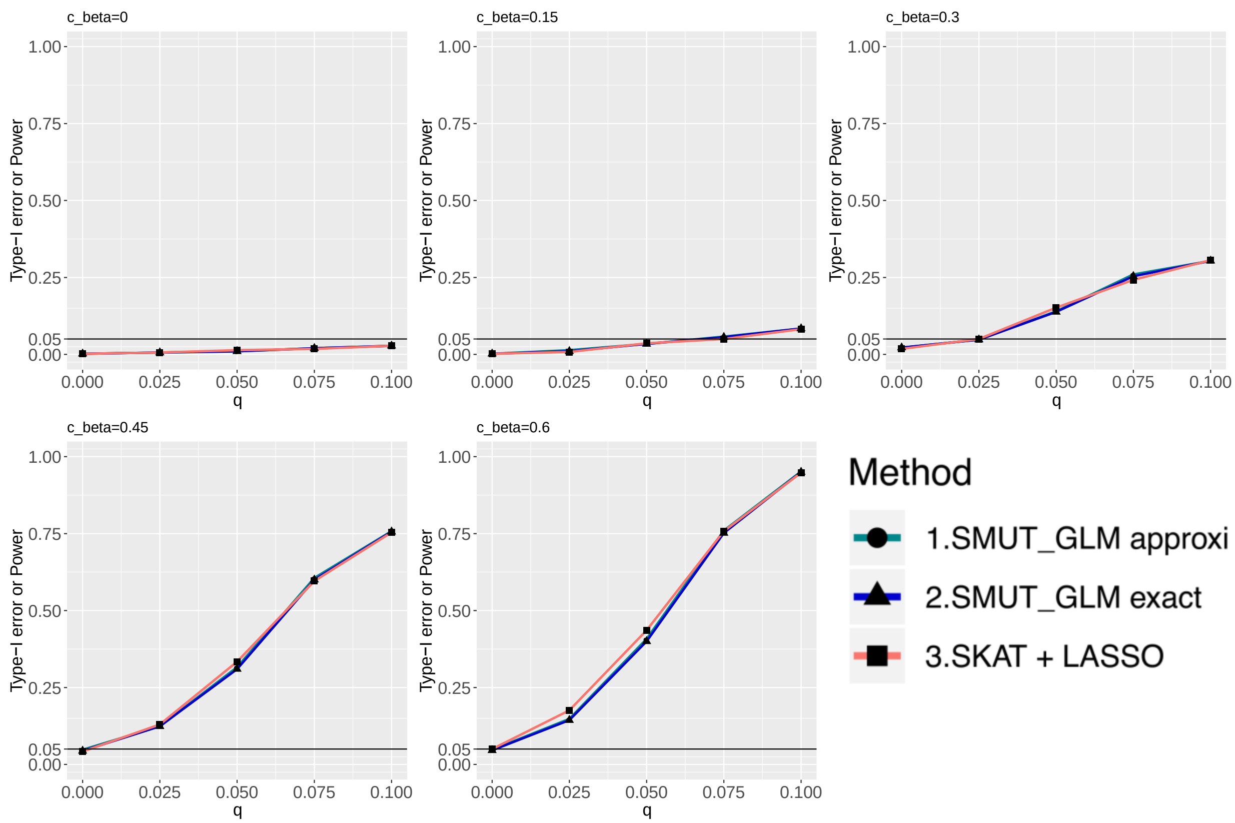


**Figure S1.** Count outcome, sparse scenario, power and Type-I error

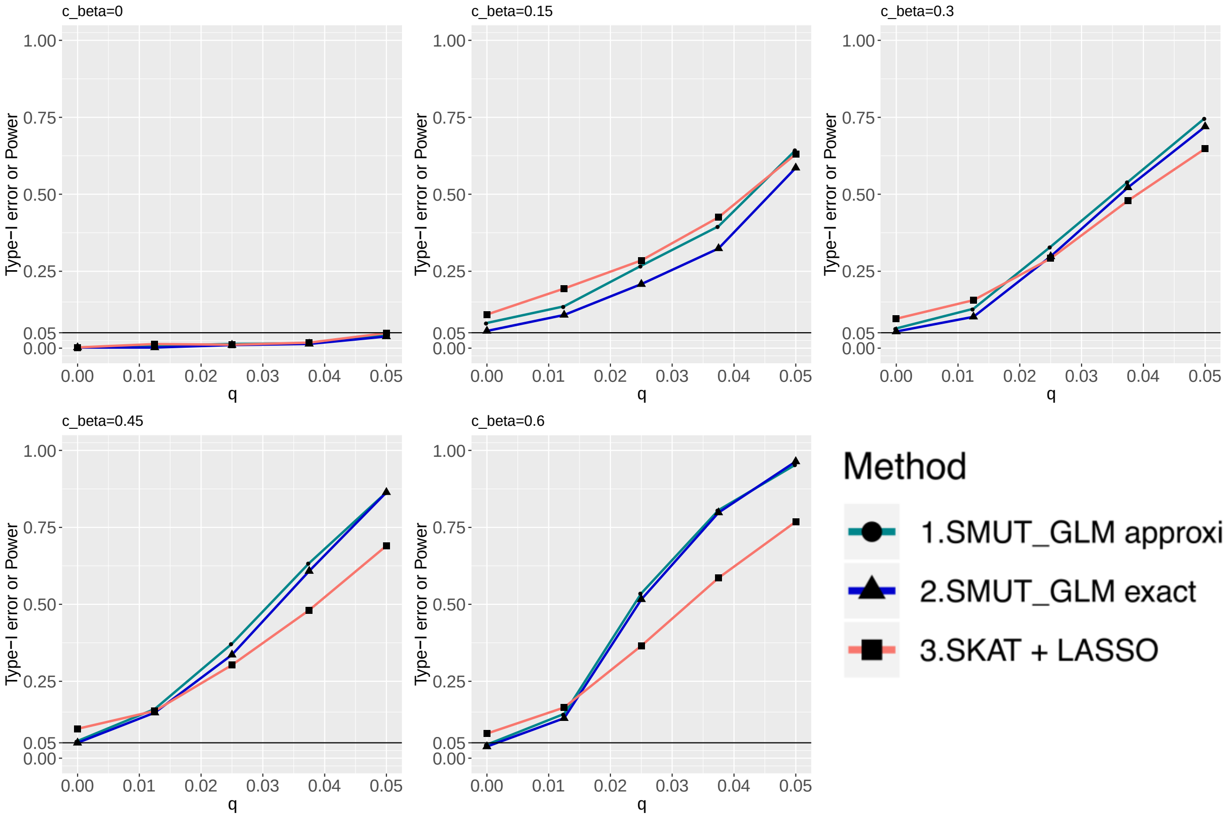


**Figure S2.** Count outcome, dense scenario, power and Type-I error (supplementary)

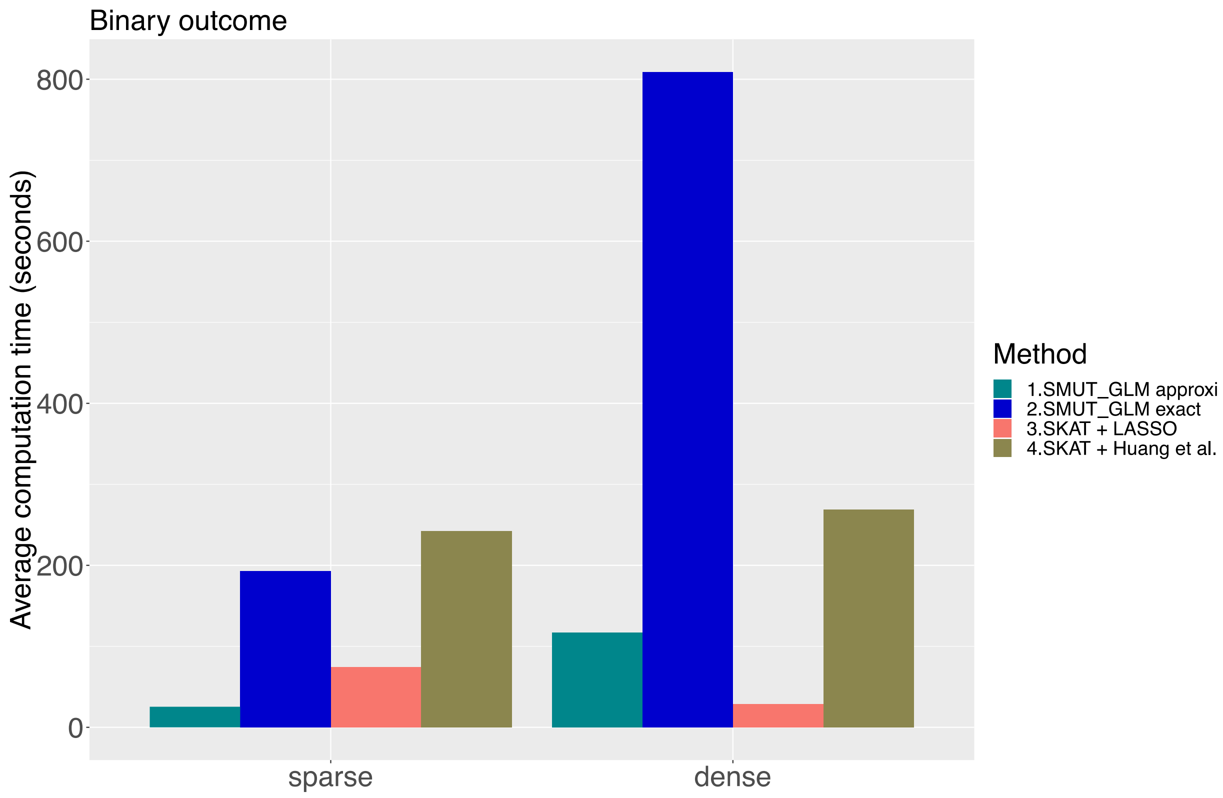


**Figure S3.** Computation time (in seconds) per dataset for binary outcome

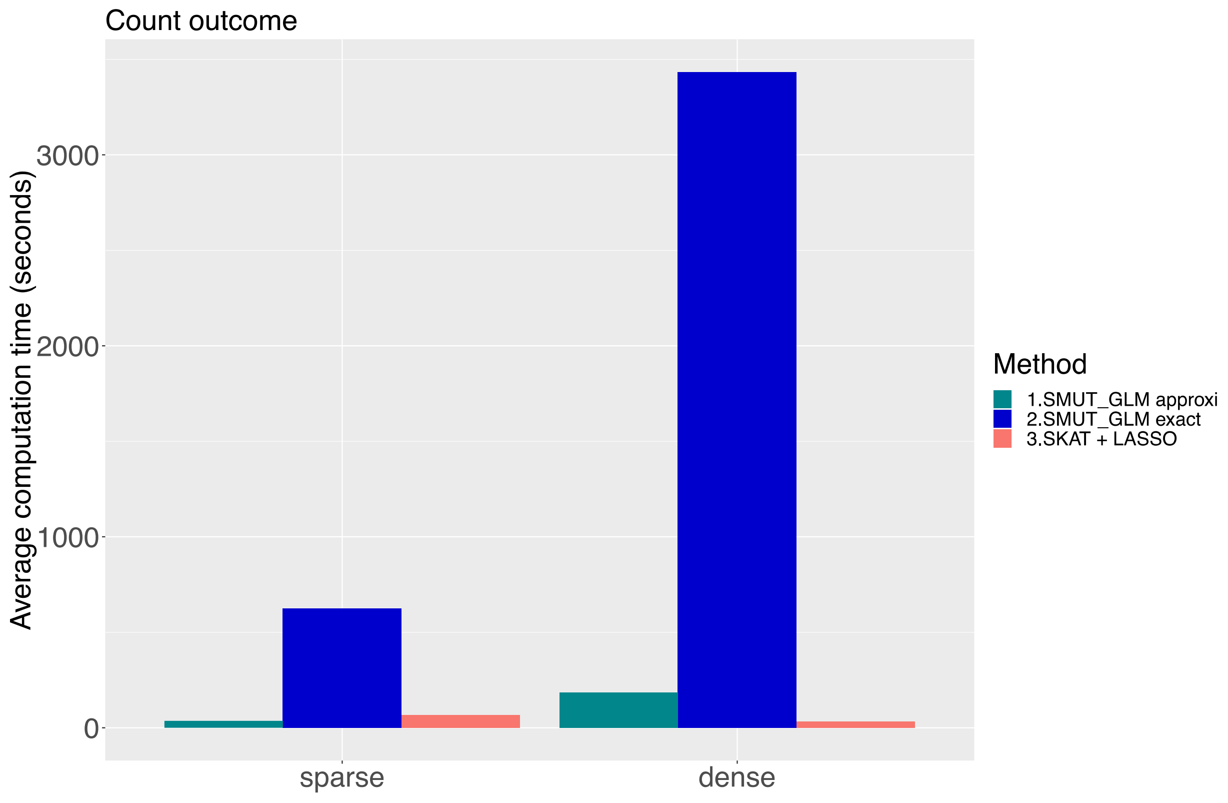


**Figure S4.** Computation time (in seconds) per dataset for count outcome

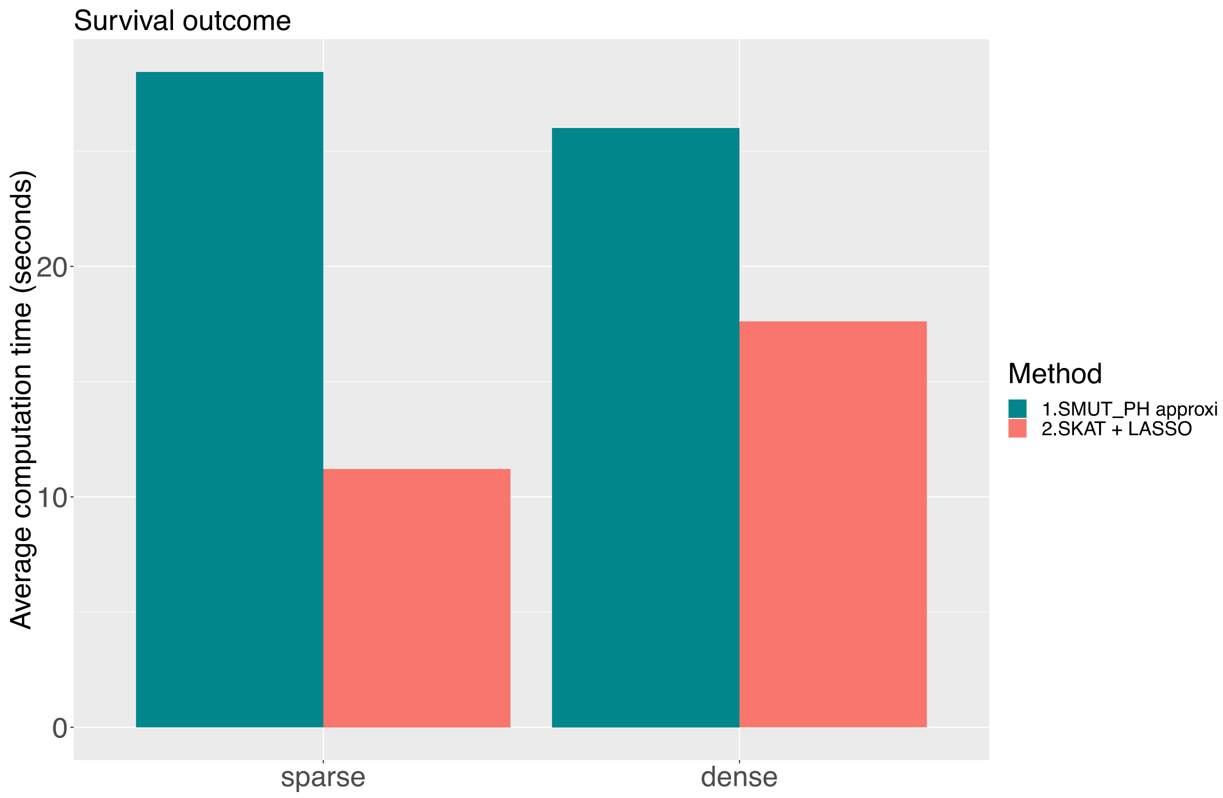


**Figure S5.** Computation time (in seconds) per dataset for survival outcome
